# A long-range chromatin interaction regulates SATB homeobox 1 gene expression in trophoblast stem cells

**DOI:** 10.1101/2020.09.11.294181

**Authors:** Wei Yu, V. Praveen Chakravarthi, Shaon Borosha, Anamika Ratri, Khyati Dalal, Michael W. Wolfe, Rebekah R. Starks, Geetu Tuteja, M.A. Karim Rumi

**Affiliations:** Department of Pathology and Laboratory Medicine, University of Kansas Medical Center, Kansas City, KS; Department of Molecular and Integrative Physiology, University of Kansas Medical Center, Kansas City, KS; Institute for Reproduction and Perinatal Research, University of Kansas Medical Center, Kansas City, KS; Department of Genetics, Development and Cell Biology, Iowa State University, Ames, IA

**Keywords:** SATB homeobox 1, trophoblast stem cells, transcriptional regulation, distant-acting enhancer, chromatin looping

## Abstract

SATB homeobox proteins are important regulators of developmental gene expression. Among the stem cell lineages determined during early embryonic development, trophoblast stem (TS) cells exhibit robust SATB expression. Both SATB1 and SATB2 act to maintain trophoblast stem-state. However, the molecular mechanisms that regulate TS-specific *Satb* expression are not yet known. We identified *Satb1* variant 2 as the predominant transcript in trophoblasts. Histone marks, and RNA polymerase II occupancy in TS cells indicated active state of the promoter. A novel cis-regulatory region with active histone marks was identified ∼21kbp upstream of variant 2 promoter. CRISPR/Cas9 mediated disruption of this sequence decreased *Satb1* expression in TS cells and chromatin conformation capture confirmed looping of this regulatory region into the promoter. Scanning position weight matrices across the enhancer predicted two ELF5 binding sites in close vicinity of SATB1 sites, which were confirmed by chromatin immunoprecipitation. Knockdown of ELF5 downregulated *Satb1* expression in TS cells and overexpression of ELF5 increased the enhancer-reporter activity. Interestingly, ELF5 interacts with SATB1 in TS cells, and the enhancer activity was upregulated following SATB overexpression. Our findings indicate that trophoblast-specific *Satb1* expression is regulated by long-range chromatin looping of an enhancer that interacts with ELF5 and SATB proteins.

## INTRODUCTION

SATB homeobox proteins (SATB1 and SATB2) are global chromatin organizers and transcriptional regulators important for tissue specific gene expression and cell lineage development. SATB proteins bind to AT-rich elements in matrix-attachment regions of actively transcribing DNA and interact with chromatin remodeling proteins as well as transcription factors to activate or repress gene expression [1-6]. SATB proteins play key roles in developmental processes, such as T cell differentiation [7-9], erythroid development [10], osteoblast differentiation and craniofacial patterning [11], cortical neuron organization [12-14], hematopoietic stem cell self-renewal [15], and embryonic stem (ES) cell pluripotency [16]. A recent study has reported that SATB proteins play distinct roles in lineage determination during early embryonic development [17]. In our previous studies, we demonstrated that SATB proteins act to maintain the trophoblast cell stem-state and inhibit trophoblast differentiation [18, 19].

SATB proteins are expressed abundantly in both mouse and rat trophoblast stem (TS) cells while in the stem-state, but the expression declines during differentiation [18, 19]. During early gestation, trophoblast cells also show high levels of SATB expression, which decreases with the progression of gestation [18, 19]. Differential expression in the trophoblast stem-state indicates a potential role for TS-specific transcriptional regulators in controlling *Satb1* expression. However, the mechanisms responsible for regulating *Satb1* gene expression in TS cells or in the placenta are currently unknown.

SATB proteins are important regulators of TS cell renewal and differentiation [19]. TS cells are the precursors of specialized differentiated cell types in the placenta. Self-renewal of TS cells and regulated differentiation into multiple trophoblast lineages are essential for proper placental development, function and maintenance of pregnancy [20-22]. SATB proteins are a part of a regulatory network that controls the development of the trophoblast lineage and regulates their differentiation. Insight into the transcriptional regulation of SATB expression in trophoblast cells will provide opportunities to manipulate its expression, which could have a wide range of applications in experimental biology.

In this study, we detected *Satb1* transcript variants expressed in trophoblast cells, and determined their promoters. We also identified a distant-acting cis enhancer that forms a long-range chromatin interaction with the proximal promoter to regulate trophoblast-specific *Satb1* expression.

## MATERIALS AND METHODS

### Cell culture

Two TS cell models were included in this study: mouse TS cells and Rcho1 rat TS cells. Mouse TS cells (obtained from Dr. Janet Rossant, Hospital for Sick Children, Toronto, Canada) were maintained in FGF4/ heparin supplemented TS culture medium [containing 30% TS basal medium (RPMI supplemented with 20% FBS, 1mM sodium pyruvate and 100μM 2-mercaptoethanol), 70% mouse embryonic fibroblast-conditioned medium, 25ng/ml FGF4 and 1μg/ml heparin] as described previously [23]. Differentiation of the cells was induced by removal of FGF4, heparin and mouse embryonic fibroblast conditioned medium [23]. ES-E14Tg2A (E14) mouse embryonic stem (ES) cells (obtained from ATCC, Manassas, VA) were maintained in RESGRO (SCM001) culture media (EMD Millipore) on feeder-free, gelatin-coated culture dishes. Extraembryonic endoderm stem (XEN) cells (obtained from Dr. Janet Rossant) were grown in Base XEN medium (RPMI supplemented with 15% FBS, 1 mM sodium pyruvate and 50μM 2-mercaptoethanol) as published earlier [24]. Rcho-1 TS cells (a rat choriocarcinoma cell line obtained from Dr. Michael Soares, University of Kansas Medical Center, Kansas City, KS) were maintained in TS basal medium (RPMI supplemented with 20% FBS, 1mM sodium pyruvate and 50μM 2-mercaptoethanol), as previously reported [25]. Differentiation was induced by growing the cells to near confluence and removing FBS [25]. 293FT cells (purchased from Thermo Fisher Scientific) were maintained in DMEM supplemented with 10% FBS and 4mM glutamine. All cell cultures were carried out at 37°C in a humidified 5% CO2 atmosphere.

To reprogram ES cells, pCAG-hCdx2ERT2-ires-puro (obtained from Dr. Jon Draper, McMaster University, Canada) or pCAG-hGata3ERT2-ires-puro (obtained from Dr. Janet Rossant) vectors were stably transfected into E14 mouse ES cells using lipofectamine 2000 (Thermo Fisher scientific). Cells were selected for puromycin resistance, and transgenes were activated by supplementing TS medium with 1 μg/ml 4-OH tamoxifen (Millipore Sigma). Cells were fed daily with the tamoxifen containing TS medium for 6 days and analyzed for gene expression [26]. Human ES cells H9 (WA09, WiCell Research Institute, Inc) were converted to trophoblasts by exposing them to BMP4, A83-01 and PD173074 in the absence of FGF2 for 2 days and analyzed for gene expression [27].

### Gene expression analysis

Gene expression analysis at the mRNA level was performed by conventional RT-PCR, RT-qPCR and RNA-seq, whereas cellular protein expression was assessed by immunofluorescence and western blot analysis.

#### RT-PCR and qRT-PCR

RNA was extracted by using TRI Reagent (Sigma-Aldrich) according to manufacturer’s instructions. cDNAs were reverse transcribed from 2μg of total RNA by using Applied Biosystems High-Capacity cDNA Reverse Transcription Kits (Thermo Fisher Scientific). Conventional PCR amplification of cDNA was done in a 25μl reaction volume by using DreamTaq Green DNA polymerase (Thermo Fisher Scientific). Real-time RT-qPCR amplification of cDNAs was carried out in a 20μl reaction mixture containing Applied Biosystems Power SYBR Green PCR Master Mix (Thermo Fisher Scientific). Amplification and fluorescence detection of qRT-PCR were carried out on Applied Biosystems StepOne Real Time PCR System (Thermo Fisher Scientific). The ΔΔCT method was used for relative quantification of target mRNA normalized to 18S RNA. All PCR primers were designed using Primer3 [28] and the sequences are shown in Table S1-S3.

#### RNA sequencing

RNA-Seq data was previously generated and analyzed [29]. FPKM values were extracted from data deposited in GEO, under accession GSE65808.

#### Immunofluorescent Microscopy

Mouse ES, TS or XEN cells were grown on coverslips placed in six-well tissue culture plates. After fixation in 4% formaldehyde for 10 min and permeabilization in 0.5% Triton X-100 for 10 min, the coverslips were blocked with 5% BSA for 1h at room temperature. After blocking, the cells were incubated with appropriately diluted primary antibodies: anti-SATB1 (ab109122, Abcam at 1:1000) and either anti-CDX2 (cdx2-88, BioGenex at 1:200), or anti-OCT4 (Sc-5279, Santa Cruz Biotechnology at 1:200) or anti-GATA4 (sc-25310, Santa Cruz Biotechnology at 1:200) at room temperature for 2h. After washing the unbound primary antibodies, secondary antibody staining was performed with Alexa Fluor 568- or 488-labeled detection reagents (goat anti-rabbit, goat anti-mouse antibodies; Molecular Probes) at 1:200 dilution, and DNA staining was performed by DAPI (Prolong Gold Antifade Mountant, Thermo Fisher Scientific). The images were captured on a Nikon Eclipse 80i microscope.

#### Western Blotting

Cell lysates were prepared in 1x SDS Sample Buffer (62.5 mM Tris-HCl pH 6.8, 2%SDS, 42mM DTT, 10% glycerol and 0.01% bromophenol blue; Cell Signaling Technology), sonicated to shear DNA and reduce viscosity and then heat denatured. Proteins were separated on 4-20% SDS-PAGE and transferred to PVDF membranes. Membranes were blocked with 5% milk and incubated with primary antibodies for 1h at room temperature. Then the membranes were incubated with following primary antibodies at appropriate dilution in blocking buffer: ant-SATB1 (ab109122, Abcam 1: 10000), anti-SATB2 (ab92446, sc-81376, 1:2000), anti-CDX2 (Abcam, 1:5000), anti-OCT4 (sc-5279, Santa Cruz Biotechnology, 1:2000), anti-GATA4 (sc-25310, Santa Cruz Biotechnology, 1:2000), anti-FLAG (#14793, Cell Signaling Technology, 1:5000) and ELF5 (sc-9645, Santa Cruz Biotechnology, 1:2000). Anti-TUBA (MABT522, Millipore Sigma, 1:20000), anti-ACTB (A5441, Millipore Sigma, 1:30000) or anti-Histone H3 (ab1791, Abcam, 1:20000) antibodies were used detect the expression of housekeeping genes as loading controls. Membranes were washed, blocked and incubated with peroxidase-conjugated anti-mouse, anti-rabbit or anti-goat secondary antibodies (Santa Cruz Biotechnology) at a dilution of 1:5000-20000, and immunoreactive signals were visualized using Luminata Crescendo Western HRP substrate (Millipore Sigma).

### Analysis of transcriptional landscape in *Satb1* promoter and enhancer

Trophoblast-specific *Satb1* promoters were initially located by variant specific RT-PCR and RNA sequencing as described above. The locations of the proximal promoters and the distant-acting *Satb1* enhancer were identified by analyses of H3K27ac ChIP-seq data. Identified promoters and the enhancer were further characterized for relevant histone marks and transcription factor binding by ChIP analyses.

#### ChIP-Seq analyses for H3K27ac in mouse early placentas

ChIP-Seq data was previously generated and analyzed [29]. Peak data was downloaded from GEO (GSE65807). Normalized wiggle signal tracks were generated using the bam_to_bigwig function in pybedtools [30].

#### Chromatin Immunoprecipitation (ChIP) of mouse TS and Rcho1 rat TS cells

Each ChIP sample was prepared with 15-20 million mouse TS or Rcho1 rat TS cells as described earlier [31]. Briefly, cells were cross-linked in 1% formaldehyde for 10 minutes at room temperature, quenched in 0.125M glycine for 5 minutes, washed twice with cold PBS with 0.5% IGEPAL CA-630 and resuspended in cold lysis buffer (50mM Tris-HCl, pH 8, 10mM EDTA, 0.2% SDS) in the presence of PMSF and protease inhibitor cocktail (Sigma-Aldrich) for 30 minutes. Cell lysates were diluted 1:1 with dilution buffer (0.01% SDS, 1.1% Triton X-100,1.2mM EDTA, 16.7mM Tris-HCl, pH 8.1, 167mM NaCl) then sonicated for 40 cycles (20 seconds on/60 sec off) at 70% amplitude to produce an average fragment size range of 300-600bp. Immunoprecipitation was performed using ∼2.5-5µg antibody (anti-H3K27ac: 05-1334 Millipore Sigma, anti-H3K9ac: 07-352 Millipore Sigma, anti-H3K4me3: 07-473 Millipore Sigma, anti-SATB1: ab109122 Abcam, anti-SATB2: sc-81376 Santa Cruz Biotechnology, anti-ELF5: sc-9645x Santa Cruz Biotechnology, anti-Pol II: sc-47701 Santa Cruz Biotechnology, anti-FLAG M8823 Millipore Sigma) conjugated to 50µl protein A/G magnetic beads (Dynabeads, Thermo Fisher Scientific) overnight. Bead-chromatin complexes were washed using High Salt Buffer (0.1% SDS, 1%Triton X-100, 2mM EDTA, 20mM Tris-HCl, pH 8.1, 500mM NaCl), Low Salt Buffer (0.1% SDS, 1% Triton X-100, 2mM EDTA, 20mM Tris-HCl, pH 8.1, 150mM NaCl), LiCl Buffer (0.25M LiCl, 1% IGEPAL, 1% Deoxycholic acid, 1mM EDTA, 10mM Tris-HCl, pH 8.1) and TE buffer (10mM Tris-HCl, 1mM EDTA, pH 8.0), with each wash performed twice for 5 minutes. Cell lysis, sonication, immunoprecipitation and cleanup steps were all performed at 4 °C. Finally, chromatin DNA was eluted from the magnetic beads using elution buffer (1% SDS, 0.1M NaHCO3), protein-DNA crosslinks were reversed with the addition of 5M NaCl and heating on a shaker incubator overnight and purified using Qiaquick columns (Qiagen). DNA was eluted in 100µl of 10mM Tris-HCl and 2.5 to 5 µl aliquots were used in qPCR analyses. qPCR primers for the target sites are shown in Table S4. Mouse positive control primer set Actb2 (#71017, Active Motif) and mouse negative control primer set 1 (#71011, Active Motif) were used for validating the ChIP assays (Fig. S1).

##### Characterization of the distant-acting Satb1 enhancer

Requirement of the distant-acting enhancer in transcriptional regulation of *Satb1* was assessed by targeted disruption of the locus using CRISPR/Cas9. Chromatin looping and interaction of the distant enhancer with the proximal promoter was demonstrated by Chromatin Conformation Capture (3C).

#### CRISPR/Cas9 mediated interference and deletion of the enhancer

CRISPR guide RNAs that specifically target the *Satb1* var2 promoter and *enhancer S* were designed to have limited off-targets using an online tool (http://crispr.mit.edu/). All gRNA sequences are listed in Table S5. Oligonucleotides encoding the gRNAs were annealed and cloned into the phU6-gRNA (Addgene, Plasmid #53188) [32] following guidelines from the Zhang lab (http://www.genome-engineering.org/crispr/?page_id=23). Rcho1 TS cells, a commonly used rat TS cell model, was selected for the CRISPR/Cas9 mediated targeted deletion experiments because of its high transfection efficiency. For CRISPR/Cas9 mediated targeted deletion of the enhancer, Rcho1 cells were stably cotransfected with the vectors (phU6-gRNA) expressing enhancer gRNAs and Cas9 (pLV hUbc-Cas9-T2A-GFP, Addgene, Plasmid #53190)[32] using Lipofectamine 2000 transfection reagent (ThermoFisher Scientific) and selected for G418 resistance and GFP expression. Selected cells were screened for targeted deletion of *Satb1* enhancer (Δ Enh S) using the PCR primers in Table S6 and characterized for trophoblast stem and differentiation markers. For CRISPR-interference, Rcho1 cells were co-transfected with the gRNA and dCas9 expression vector (pLV hUbc-dCas9-T2A-GFP; Addgene, Plasmid #53191) [26]. After 3 days of transfection, cells were harvested for RNA isolation and analyses of *Satb1* expression.

#### Chromatin Conformation Capture (3C)

3C was carried out following a standard protocol [33]. 3C experiments performed in mouse TS cells were compared with that in mouse embryonic fibroblasts that do not express *Satb1*. Briefly, mouse TS cells and mouse embryonic fibroblasts were fixed in 1% formaldehyde for 10 min at room temperature. After quenching the crosslinking reaction with 0.125 M glycine for 5 min, cells were washed with cold PBS, resuspended in cold lysis buffer (10mM Tris-HCl pH 7.5, 10 mM NaCl, 5 mM MgCl2, 0.1 mM EGTA with protease inhibitors) and incubated on ice for 30 min. After centrifugation at 2000g for 5 min, pelleted nuclei were resuspended in 2 ml of cold lysis buffer. Approximately 10^7^ nuclei were resuspended in 500μl of 1.2x FastDigest Restriction Enzyme Buffer (Thermo Fisher Scientific) containing 1.6% SDS and incubated for 1 h at 37°C with shaking at 250 rpm. SDS was subsequently quenched by adjusting the reaction to 2% Triton-X100 followed by another 1h incubation at 37°C with shaking. An aliquot of 20μl was taken from each sample and stored at −20°C for use as undigested genomic DNA. Then 50μl of FastDigest *Bgl* II restriction enzyme (Thermo Fisher Scientific) was added to the reaction tube and incubated overnight at 37°C with shaking at 250rpm. The restriction enzyme was deactivated by adding 40μl of 20% SDS and heating at 65°C for 20 min. The reaction was diluted in 7ml of 1.1x T4 DNA ligase reaction buffer (Thermo Fisher Scientific), and 375μl of 20% Triton-X100 was added and incubated at 37°C for 1h to quench SDS. Digested chromatin was ligated with 150U of T4 DNA ligase (Thermo Fisher Scientific) for 4h at 16°C. Formaldehyde crosslinks were reversed with Proteinase K digestion and overnight incubation at 65°C. RNAs were degraded with RNase treatment at 37°C for 1h. 3C libraries were purified by phenol-chloroform extraction and precipitated with 2.5 volumes of 100% ethanol and 0.1 volume of 3M sodium acetate and incubating at −80°C for 1h. Precipitated DNA was collected by centrifugation at 5000g for 1h and washed in 70% ethanol. DNA pellets were resuspended in 150µl of 10 mM Tris-HCl pH 7.5 and 3C products were checked by conventional PCR. PCR primers used in 3C analysis are shown in Table S7.

##### Transcription factor binding to the distal enhancer

Putative ELF5 and SATB1 binding sites were identified in the *Satb1* enhancer (chr17: 51993298-51994604) using TFBSTools [34], and a 90% match threshold. Position weight matrices (PWMs) for ELF5 and SATB1 were obtained from a motif library described previously [35]. This analysis predicted multiple ELF5 binding sites near SATB1 binding sites. Further confirmation of these potential transcription factor binding sites was done by enhancer-reporter luciferase assays, ChIP analyses and investigating a possible interaction between ELF5 and SATB1.

#### Luciferase reporter assays

To prepare the enhancer-reporter constructs, the *Satb1* enhancer sequence was cloned into the KpnI and XhoI sites of pGL4.25[luc2CP/minP] firefly luciferase vector containing a minimal TATA promoter (Promega). Rcho1 TS cells were used for the reporter assay. Twenty-four h after plating in 12-well plates, Rcho1 cells were transfected with the enhancer-reporter vector along with a control Renilla luciferase vector (pGL4.74 [hRluc/TK]) using Lipofectamine 2000 (Thermo Fisher Scientific). Expression vectors for SATB1, SATB2 or ELF5 were individually cotransfected with the reporter vector to assess their regulatory role on the enhancer sequence. 12h after the transfection, transfection medium was replaced with cell proliferation medium and cultured for another 12h. 24h after transfection, cells were washed with cold PBS, lysed in 100µl of passive lysis buffer and standard dual luciferase assays were performed on the cell lysates by using Dual-Luciferase Reporter Assay reagents (Promega).

#### ChIP assays

ChIP assays were performed as describe above.

#### ELF5-SATB1 interaction

Protein-protein interaction was investigated by co-immunoprecipitation. Rcho1 cells stably expressing FLAG-tagged SATB1 or ELF5 were harvested to extract nuclear proteins. Nuclear proteins were extracted in nondenaturing buffer (20mM Tris-HCl pH 7.5, 2mM EDTA) adjusted to 0.3 M NaCl and 0.5% Triton X-100. After centrifugation at 40,000g for 1h at 4C in a Ti-70 rotor, the supernatants were mixed with anti-FLAG (M2) magnetic beads (Millipore Sigma) at a ratio of 100 μl of beads/1 ml of nuclear extract and gently rocked overnight at 4°C. The beads with immunoprecipitated protein complexes were washed 8 times with wash buffer containing 50mM Hepes-NaOH, pH 7.9, 0.25 M KCl, 0.1% Triton X-100, and then eluted with 200μl of wash buffer containing 0.4mg/ml FLAG peptide (Millipore Sigma). Eluted proteins were mixed with 2xSDS sample buffer, boiled for 10min, separated on SDS-PAGE, and processed for Western blot analysis.

### ELF5 regulation of *Satb1* expression in TS cells

The TS regulators ELF5 and SATB proteins demonstrated a high level of transcriptional activation of the *Satb1* enhancer in luciferase assays. We further analyzed the role of ELF5 in regulating *Satb1* expression using a ‘loss of function’ study.

#### Elf5 knockdown

For the loss of function studies, *Elf5* was knocked down in Rcho1 cells by lentiviral delivery of shRNAs. *Elf5* shRNAs, cloned into the lentiviral vector pLKO.1, were obtained from Millipore Sigma (St. Louis, MO). A control shRNA that does not target any known mammalian gene, pLKO.1-shSCR (Addgene, Plasmid #1864), was obtained from Addgene (Cambridge, MA). Lentiviral packaging vectors from Addgene (pMDLg/pRRE Plamid # 12251, pRSV-Rev Plasmid #12253 and pMD2.G Plasmid# 12259) were used to produce the viral particles in 293T cells as described earlier [36]. Culture supernatants containing lentiviral particles were harvested every 24 h for 2 days, centrifuged to remove cell debris, filtered, and applied to Rcho1 cells in culture. Transduced cells were selected for puromycin resistance. *Elf5* knockdown as well as the effect of *Elf5* knockdown on *Satb1* expression was assessed by RT-qPCR assays. Functionally active shRNA sequences are shown in Table S8.

## RESULTS

### Trophoblast-specific expression of *Satb1*

Expression of *Satb1* mRNA and protein was examined in mouse TS, ES and XEN cells. Mouse TS cells exhibited a much higher level of *Satb1* expression than ES and XEN cells (Fig. 1A-C, E, F). Expression of *Satb1* in mouse TS cells declined upon induction of trophoblast differentiation (Fig. 1D). Mouse ES or XEN cells minimally express *Satb1* in the stem-state (Fig. 1A-C, E, F); however, the expression of *Satb1* was induced when mouse ES cells were reprogrammed to a trophoblast fate by overexpression of CDX2 (Fig. 1G) or GATA3 (Fig. 1H). In addition, *Satb1* expression was also increased when human ES cells were differentiated into trophoblast cells following BMP4 treatment (Fig. 1I).

**Fig. 1.**
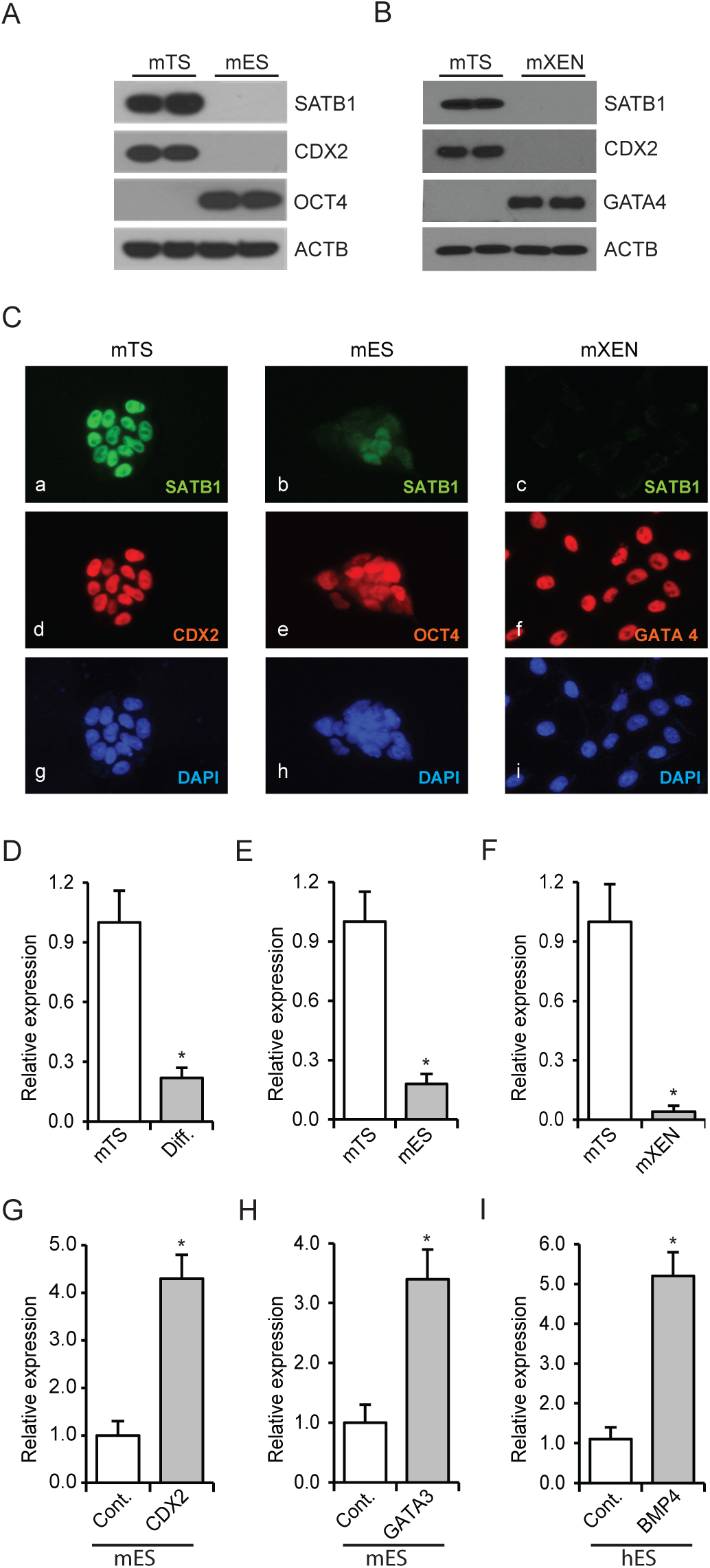
Trophoblast-specific expression of *Satb1*. Mouse trophoblast stem (mTS) cells express high levels of SATB1 compared to mouse embryonic stem (mES) cells (A) or mouse extraembryonic endoderm (mXEN) cells (B) as detected by western blotting. CDX2, OCT4 and GATA4 were detected as lineage-markers. ACTB was detected as loading control. C) Immunofluorescence imaging also detected an abundant expression of SATB1 in mTS cells (Ca). Compared to mTS cells, the level of expression is remarkably lower in mES cells (Cb) and mXEN (Cc). *Satb1* mRNA levels in TS, ES and XEN cells (D-F) correlated well with the protein expression (A-C), and in mTS cells, the mRNA level is significantly reduced upon induction of trophoblast differentiation (D). Expression of *Satb1* was induced when mES cells were reprogrammed towards trophoblast lineage by the overexpression of CDX2 (G) or GATA3 (H). BMP4 induced reprogramming of human ES (hES) cells towards trophoblast lineage also upregulated *Satb1* expression (I). RT-qPCR data are expressed as the means ± S.D. *, p< 0.05 (n=3). Diff., differentiated mTS cells; Cont., control cells.

### *Satb1* promoters in trophoblast cells

Reference sequences of four different transcript variants of mouse *Satb1* mRNA have been reported and validated (Fig. S1A, B). RT-PCR analyses suggested that the first exon in each variant is transcribed from alternative transcription start sites over a span of 21kbp of genomic DNA (Fig. 2 A-C and Fig. S1). Only variant 1 and 2 transcripts were detected in mouse trophoblast cells of e7.5 ectoplacental cones (EPCs) (Fig. 2C), with variant 2 being the predominant transcript (Fig. 2C and 3C). ChIP-sequencing (ChIP-seq) analyses for H3K27ac in mouse e7.5 EPCs demonstrated the presence of this transcription activation mark in the proximal promoters of both transcript variants (Fig. 2D). Both promoters also contained CpG islands (Fig. 2D). Next, the variant 2 promoter in mouse TS cells was examined for active histone marks. ChIP assay results supported the early placental ChIP-seq data for H3K27ac (Fig. 2E). The promoter also showed enriched marks of H3K4me3 (Fig. 2F) and RNA polymerase II (Pol II) occupancy (Fig. 2G), while the positive and negative control primer sets exhibited expected enrichment of histone marks or Pol II binding (Fig. S1).

**Fig. 2.**
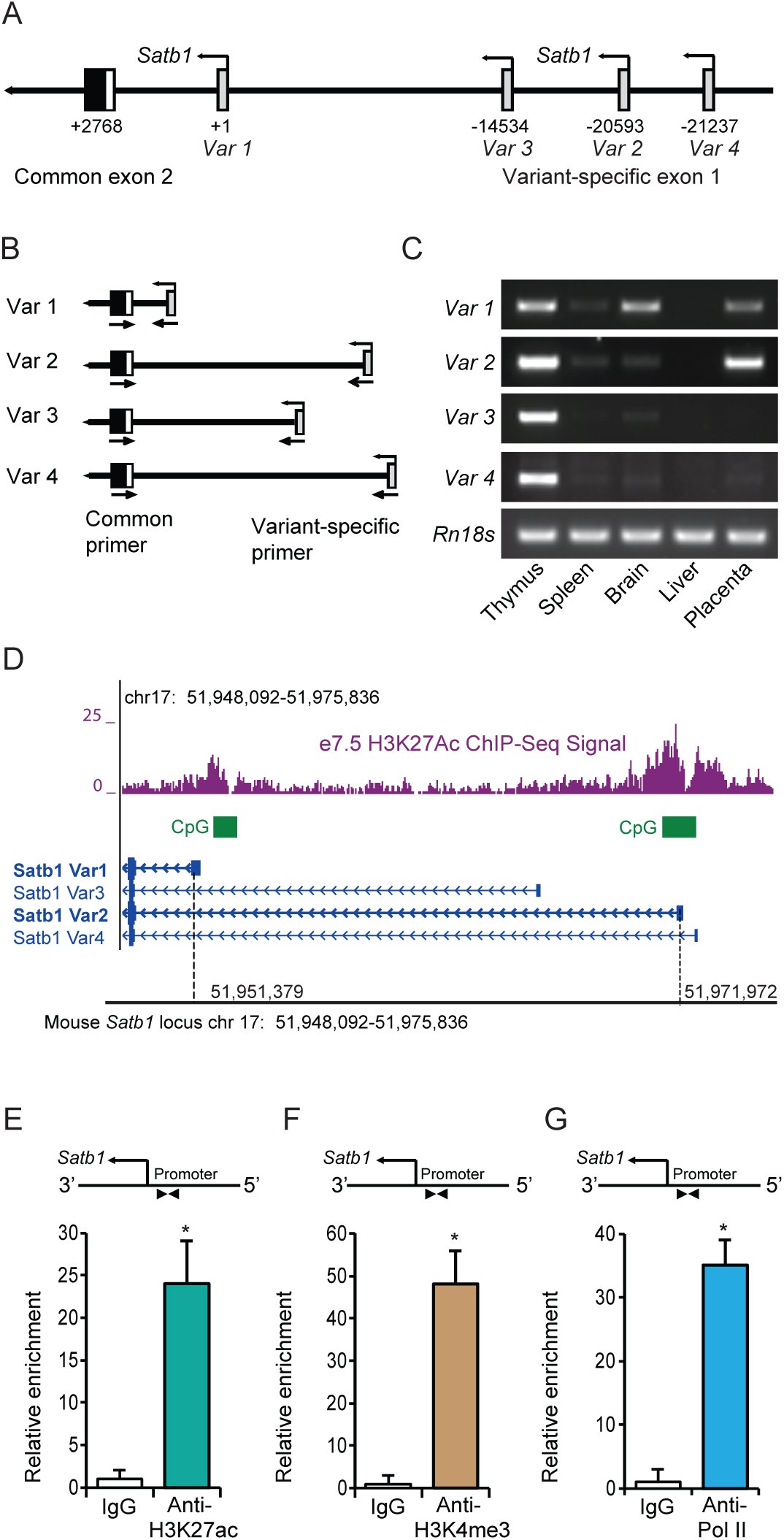
Detection of trophoblast-specific *Satb1* promoters. A) Schematic presentation of the mouse *Satb1* gene locus showing four transcript variants, each transcribed from a variant-specific alternative exon 1. Nucleotide positions are indicated with respect to the start site (TSS) of variant 1. B) Strategy of PCR-based detection of different transcript variants. C) *Satb1* transcript variants and alternative transcription start sites were detected in mouse embryonic day 7.5 (e7.5) ectoplacental cone (EPC) by RT-PCR analyses. *Satb1* transcript variants in mouse thymus, spleen, brain and liver were detected as controls for comparison. D) ChIP-seq data on e7.5 EPCs demonstrated that both variant 1 and 2 proximal promoters possessed active histone marks of acetylated histone H3 lysine 27 (H3K27ac). The promoters also contained CpG islands (D). Using ChIP assays, the variant 2 promoter in mouse TS cells was assessed for transcriptionally active histone marks of H3K27ac (E) and H3K4me3 (F), which were associated with enriched RNA polymerase II (Pol II) binding (G). ChIP-qPCR primers located in the proximal promoter region is shown schematically in E-G. The primer sequences are mentioned in Table S4. ChIP-qPCR data are expressed as the means ± S.D. *, p< 0.05 (n=3).

### Identification of a distant-acting enhancer for *Satb1* gene

RT-qPCR data indicate that the expression of both transcript variants of mouse *Satb1* was markedly reduced during differentiation of mouse TS cells *in vitro* (Fig. 3 A, B). A similar reduction in *Satb1* expression was also detected *in vivo* with RNA-sequencing (RNA-seq); expression of both variant 1 and variant 2 were significantly decreased in e9.5 compared to e7.5 placenta (Fig. 3 C). Such reductions in expression correlated well with the changes in H3K27ac activity within a potential cis-acting enhancer region (*enhancer S*) approximately 21kbp upstream of the *Satb1* variant 2 promoter (Fig. 3 D). ChIP assays using mouse TS cells also detected enriched histone marks of H3K27ac and H3K9ac, as well as enrichment of Pol II binding in the enhancer region (Fig. 3 E-G). We termed this distant-acting cis enhancer as *enhancer S*, a potential enhancer of *Satb1*.

**Fig. 3.**
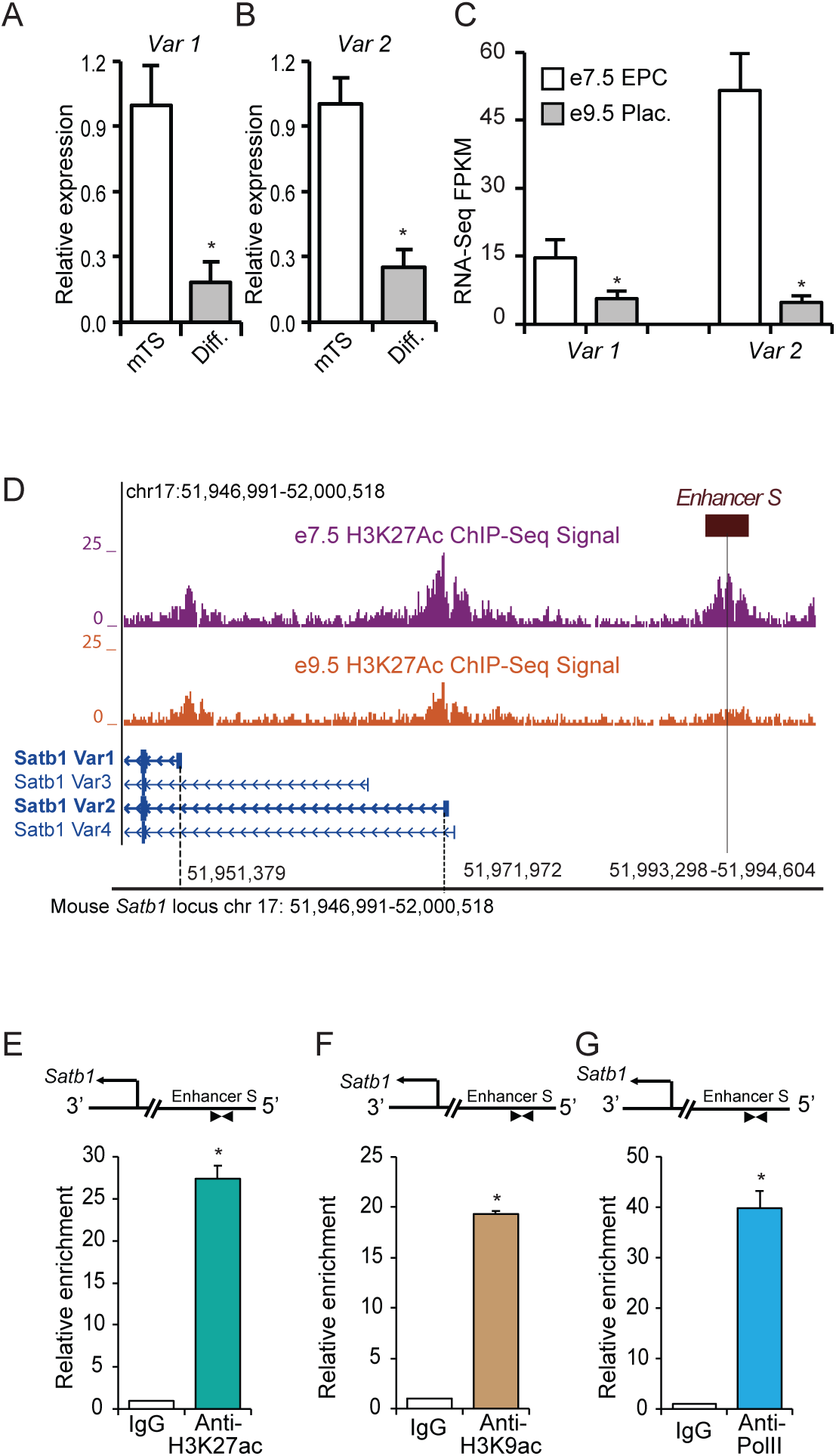
A long-distance enhancer regulates *Satb1* expression in mouse TS cells. RT-qPCR analyses indicate that expression of *Satb1* transcript variants 1 and 2 was markedly reduced in differentiated mouse TS cells (A, B). A similar reduction in *Satb1* expression was also detected by RNA-seq analyses (C). The expression of both transcript variants was markedly reduced in e9.5 mouse placenta compared to that in e7.5 placenta (C). Such reductions in *Satb1* expression level correlated with the epigenetic marks of the active chromatin state of *Satb1* promoters and an e7.5-specific distal enhancer (*enhancer S*) region ∼21kbp upstream of the variant 2 transcription start site, as detected by H3K27ac ChIP-seq (D). Mouse TS cells were positive for enrichment of H3K27ac, and H3K9ac at the potential enhancer site (E, F). Enriched Pol II binding at the enhancer was also detected by ChIP assays (G). ChIP-qPCR primers located in the enhancer region is shown schematically in F-H. The primer sequences are mentioned in Table S4. RNA-seq FPKM, RT-qPCR and ChIP-qPCR data are expressed as the means ± S.D. *, p< 0.05 (n=3). EPC, ectoplacental cone; Plac., Placenta.

### Distant-acting enhancer S is required for maintaining *Satb1* expression in TS cells

Using the CRISPR/Cas9 methodology, we investigated whether the distant enhancer was required for maintaining *Satb1* expression in TS cells. Transfection of expression vectors encoding Cas9 and the enhancer targeted gRNAs resulted in deletion of *enhancer S* in Rcho1 rat TS cells (Fig. 4 A). Deletion of the enhancer caused a dramatic reduction in *Satb1* expression (Fig. 4 B), which was associated with induction of premature differentiation in Rcho1 cells maintained in a proliferating culture condition (Fig. 4 D-H). Premature differentiation of Rcho1 cells was identified by the reduction of stem markers *Cdx2* and *Eomes*, and an increase of the differentiation marker *Prl3b1* (Fig. 4 D-F). To determine whether the reduction in *Satb1* expression was due to induction of differentiation or disruption of *enhancer S*, we further investigated its requirement using CRISPR interference. Transfection of dCas9-repressor (dCas9-KRAB) and gRNAs targeted to the enhancer sequence also markedly reduced *Satb1* expression (Fig. 4 I). CRISPR interference of *enhancer S* reduced *Satb1* expression in the same way as interference of the variant 2 promoter in Rcho1 TS cells (Fig. 4J).

**Fig. 4.**
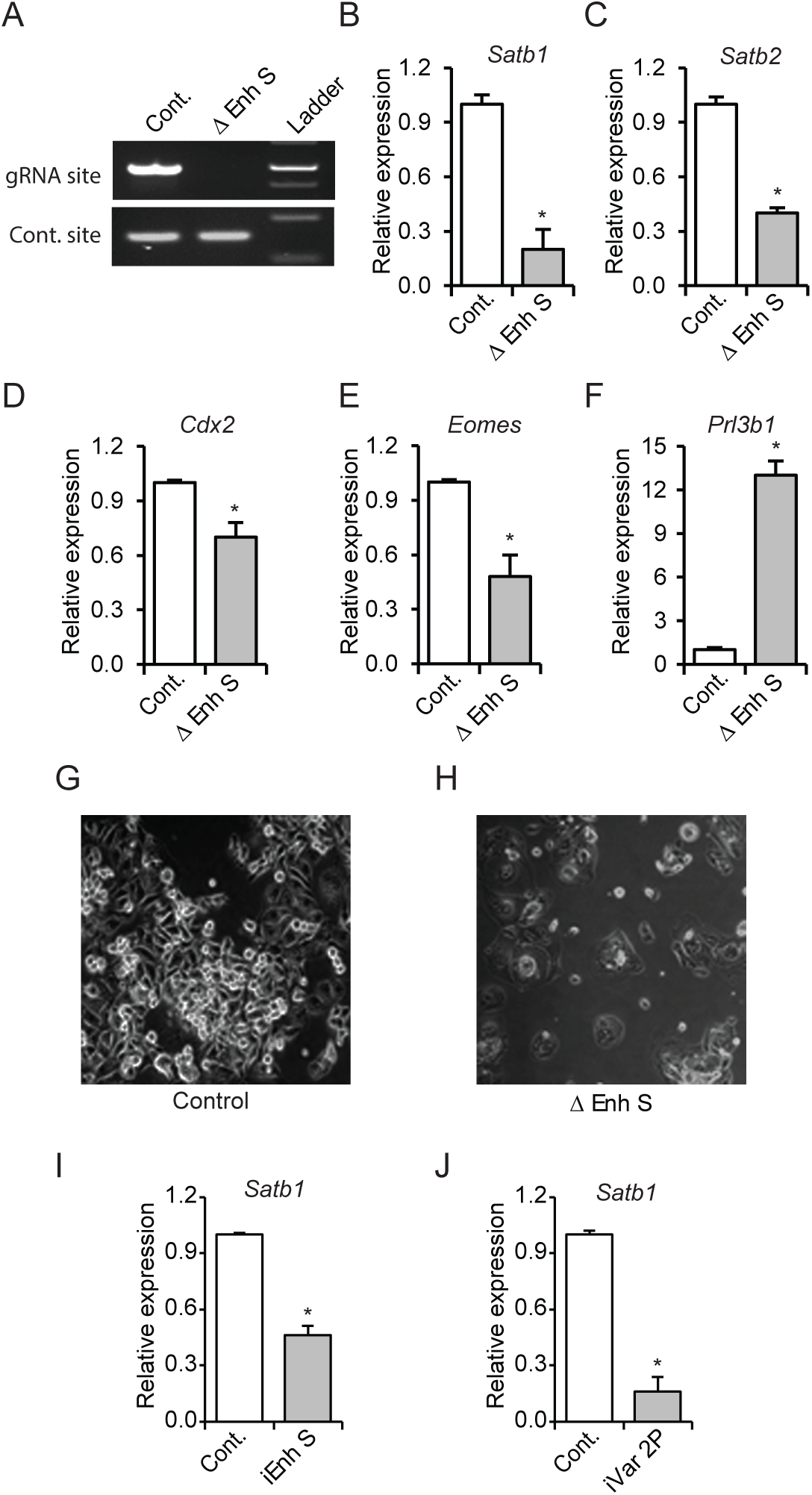
Enhancer S is required for *Satb1* expression. Rcho1 rat trophoblast cells were transfected with Cas9 and control or targeted gRNA expression constructs. Stably transfected cells were selected and assessed for targeted deletion of the enhancer. Applying the CRISPR/Cas9 system resulted in the deletion of the gRNA targeted site in *enhancer S* (Δ Enh S) (A), decreased *Satb1* expression (B) and caused differentiation of Rcho1 cells (C-H). The requirement of *enhancer S* was further confirmed by transient transfection of dCas9-KRAB and the enhancer targeted gRNAs (iEnh S) (I). Transfection of gRNAs targeted to the variant 2 promoter (iVar2P) was used as positive control (J). RT-qPCR data are expressed as the means ± S.D. *, p< 0.05 (n=3).

### Enhancer S loops into the proximal promoter to regulate *Satb1* expression

We examined the molecular mechanism as to how the distant-acting *enhancer S* regulated *Satb1* expression. Involvement of chromatin looping that can bring the enhancer into proximity with the promoter was tested by chromatin conformation capture (3C) in mouse TS cells (Fig. 5 A, B). A looping interaction between *enhancer S* and the *Satb1* variant 2 promoter was detected by 3C-PCR in mouse TS cells, but not in MEFs (Fig. 5C). Restriction analyses (Fig. 5D) and DNA sequencing (Fig. 5 E) confirmed that the 3C-PCR captured and amplified a ligation between the distant-acting *enhancer S* and the *Satb1* variant 2 promoter.

**Fig. 5.**
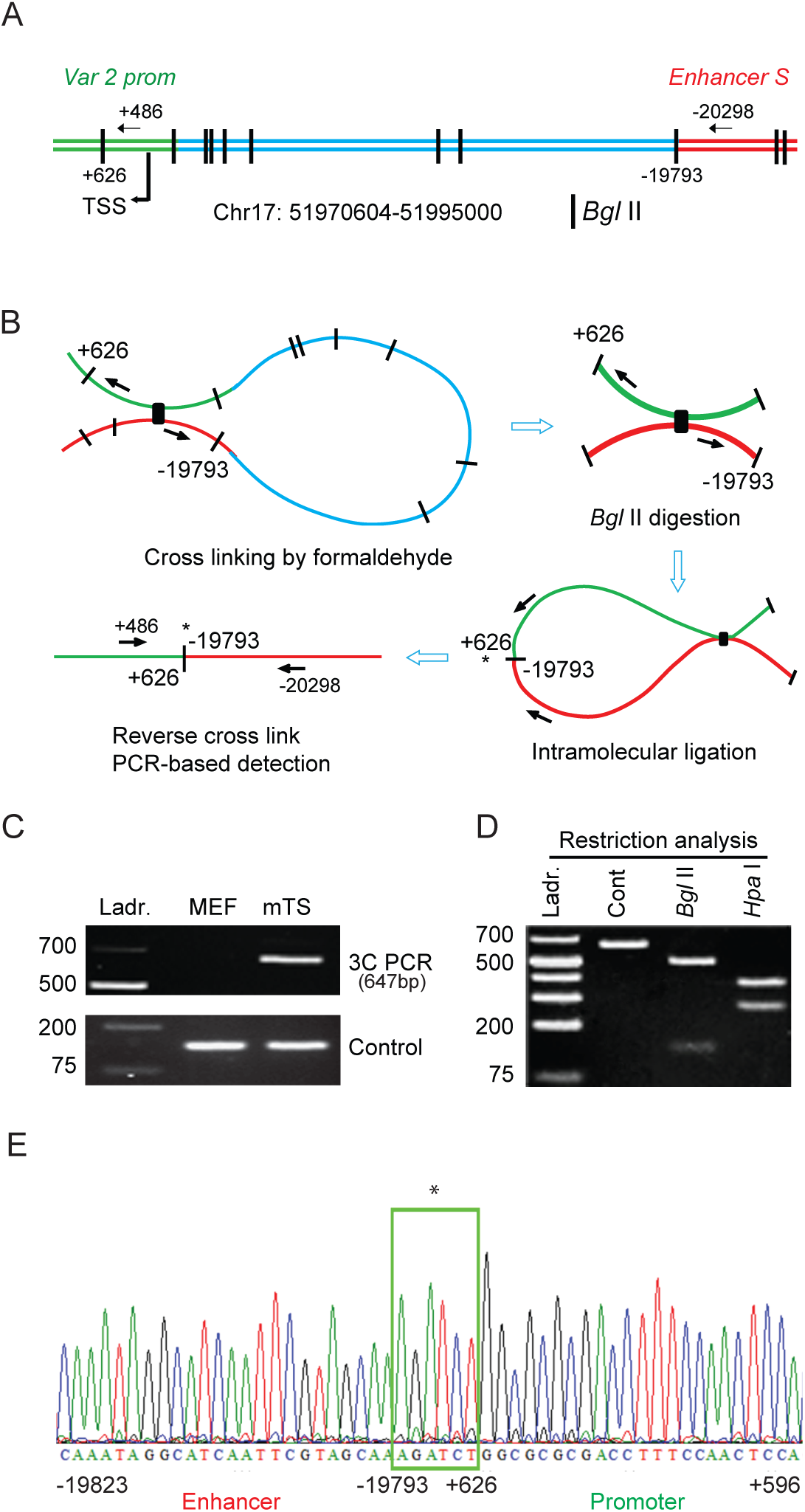
Enhancer S loops into the *Satb1* promoter in mouse TS cells. (A) Schematic diagram of the mouse *Satb1* locus showing the variant 2 promoter (var 2 prom), transcription start site (TSS), *Bgl* II restriction sites, and 3C PCR primer positions. B) Representation of the major steps of 3C PCR-based detection of the looping and interaction of *enhancer S* with the *Satb1* var 2 promoter. 3C PCR detected a physical interaction of the enhancer with the *Satb1* promoter in mouse TS (mTS) cells but not in mouse embryonic fibroblast (MEF) cells (C). The 3C PCR product (648bp) was confirmed by restriction analyses (D) as well as DNA sequencing (E). * indicates DNA ligation site. Ladr., DNA ladder.

### Transcriptional regulation of enhancer S in TS cells

Scanning position weight matrix (PWM) analyses (Fig. 6 A, B) of *enhancer S* using TFBSTools predicted two ELF5 binding sites in close vicinity of SATB1 binding sites (Fig. 6C). ChIP assays also demonstrated a marked enrichment of ELF5, SATB1 and SATB2 binding to the enhancer locus (Fig. 6 D-F) as well as the *Satb1* variant 2 promoter (Fig. 6 G-I).

**Fig. 6.**
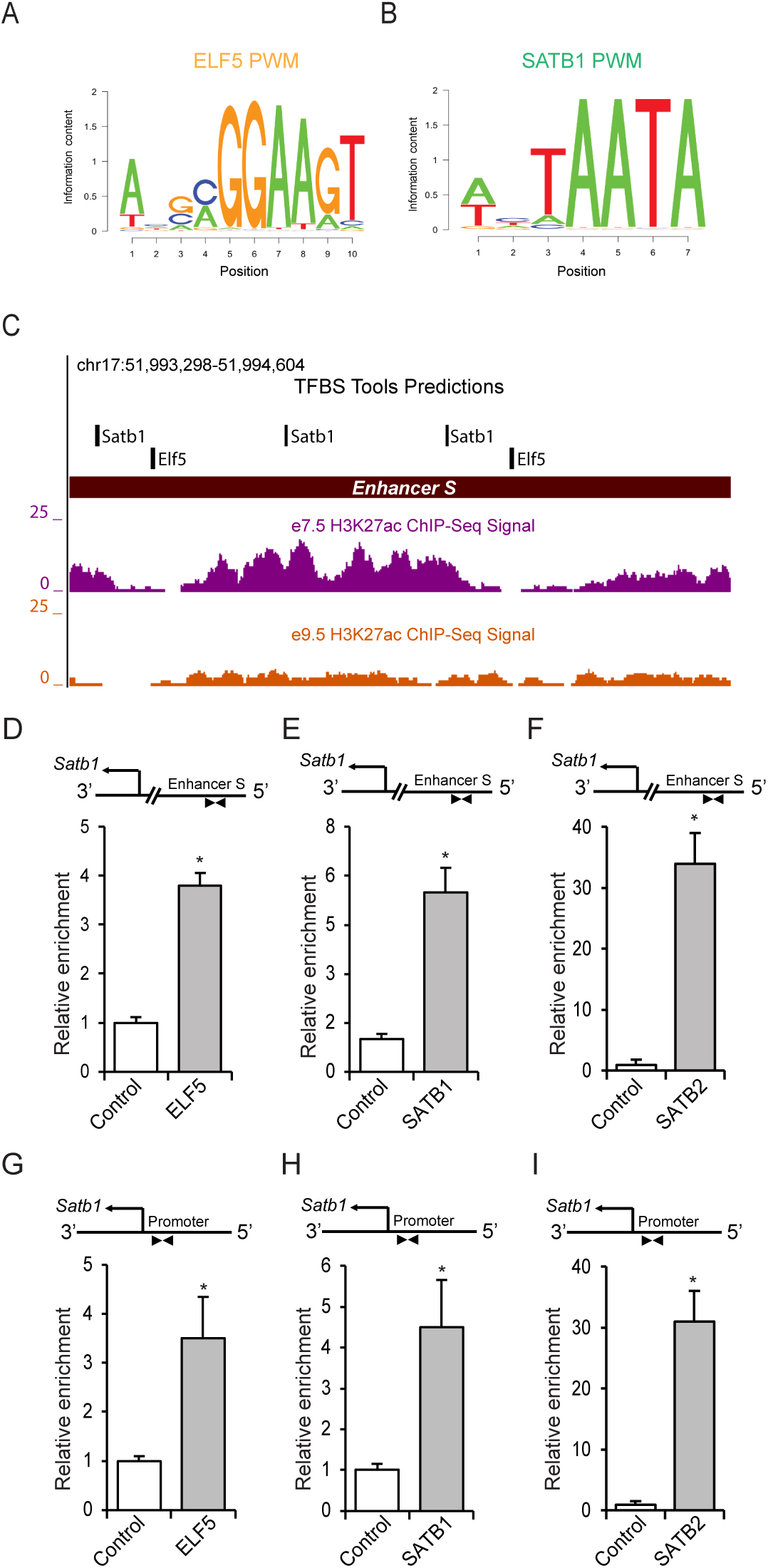
ELF5 and SATB proteins bind within the enhancer S in mouse TS cells. A, B) PWMs of ELF5 and SATB1 used for scanning the enhancer S sequence (Chr17: 51993298-51994604) using TFBSTools. C) Transcription factor binding site analysis by TFBSTools predicted the presence of two ELF5 binding sites near SATB1 binding sites within *enhancer S*. ChIP assays also demonstrated significant enrichment of ELF5, SATB1 and SATB2 in the enhancer locus of mTS cells (D-F). G-I), In addition to the enhancer region, binding of ELF5, SATB1 and SATB2 was detected in the *Satb1* variant 2 promoter in mTS cells. ChIP-qPCR data are expressed as the means ± S.D. *, p< 0.05 (n=3).

ELF5 and SATB proteins exhibited trophoblast stem-state specific differential expression both *in vivo* and *in vitro* (Fig. 7, A-F). We further analyzed the role of ELF5 in regulation of *Satb1* expression by shRNA mediated knockdown of *Elf5* in Rcho1 TS cells (Fig. 7 G, H). Knockdown of ELF5 significantly downregulated the expression of *Satb1* (Fig. 7 H).

**Fig. 7.**
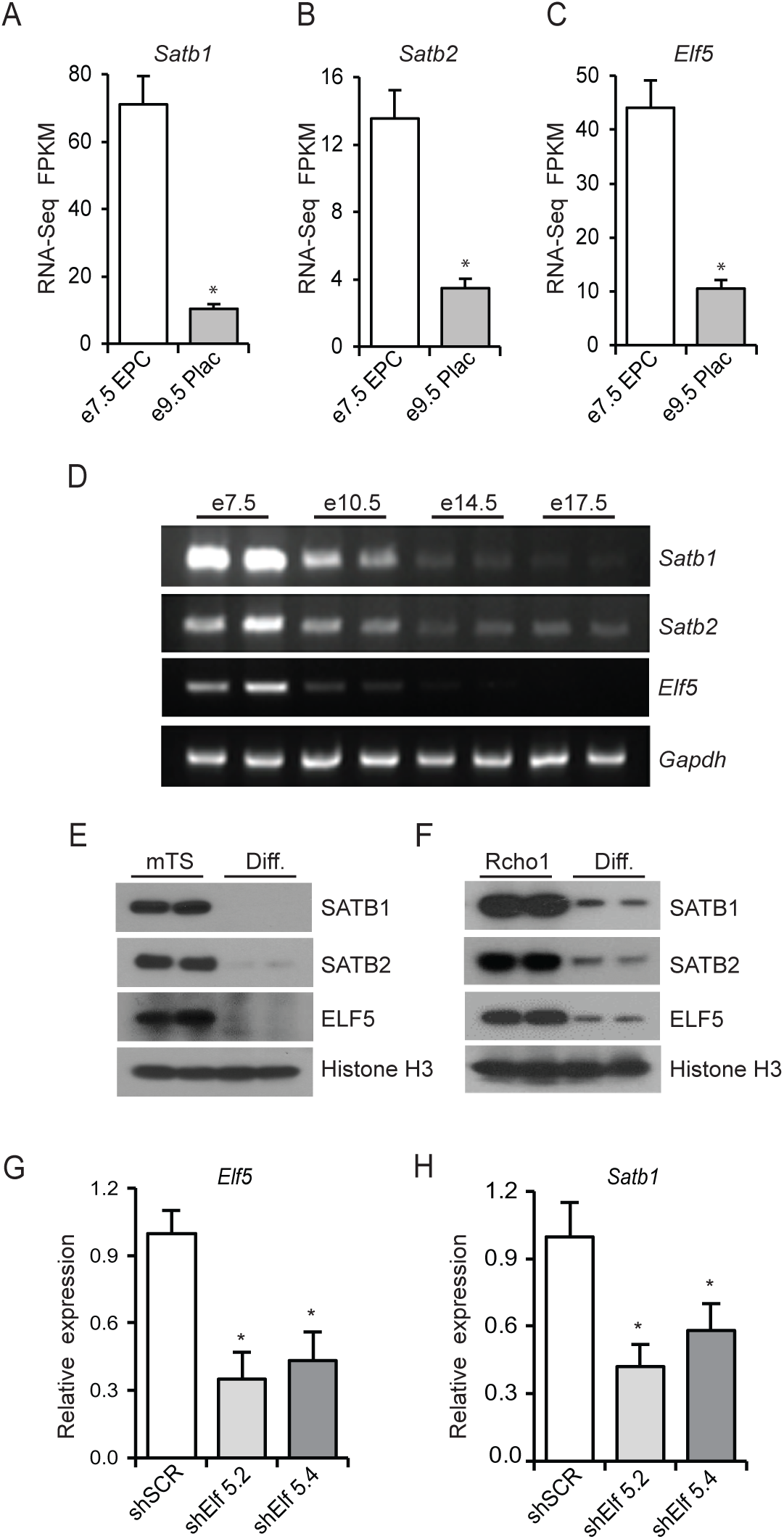
ELF5 regulates *Satb1* expression in TS cells. A-C) RNA-seq analyses show that expression of *Satb1, Satb2* and *Elf5* is dramatically reduced in mouse e9.5 placentas compared to e7.5 EPCs. Similar findings were observed by RT-PCR analyses of mouse placenta samples collected during the progression of gestation (D). Both mouse TS cells and Rcho1 rat TS cells exhibited a similar reduction in SATB1, SATB2 and ELF5 proteins with induction of differentiation (E, F). G, H), Rcho1 rat TS cells were stably transduced with *Elf5* shRNAs. shRNA mediated knockdown of *Elf5* (G) significantly reduced the *Satb1* mRNA level (H) highlighting its role in transcriptional regulation of *Satb1*. RNA-Seq FPKM and RT-qPCR data are expressed as the means ± S.D. *, p< 0.05 (n=3).

To assess the role of these transcriptional regulators on *enhancer S*, a reporter construct was prepared by cloning the enhancer upstream of a minimal TATA promoter within pGL4.25[luc2CP/minP] firefly luciferase vector (Fig. 8B). Cotransfection of the enhancer-reporter and expression vectors for ELF5, SATB1 or SATB2 into Rcho1 rat TS cells significantly upregulated reporter activity (Fig. 8 C-E). Furthermore, co-immunoprecipitation of either ELF5 or SATB1 with Rcho1 nuclear proteins detected an interaction between ELF5 and SATB1 (Fig. 8 F, G). Taken together, we propose a model of ELF5-SATB1 interaction that regulates *Satb1* expression in the trophoblast stem-state (Fig. 8 H).

**Fig. 8.**
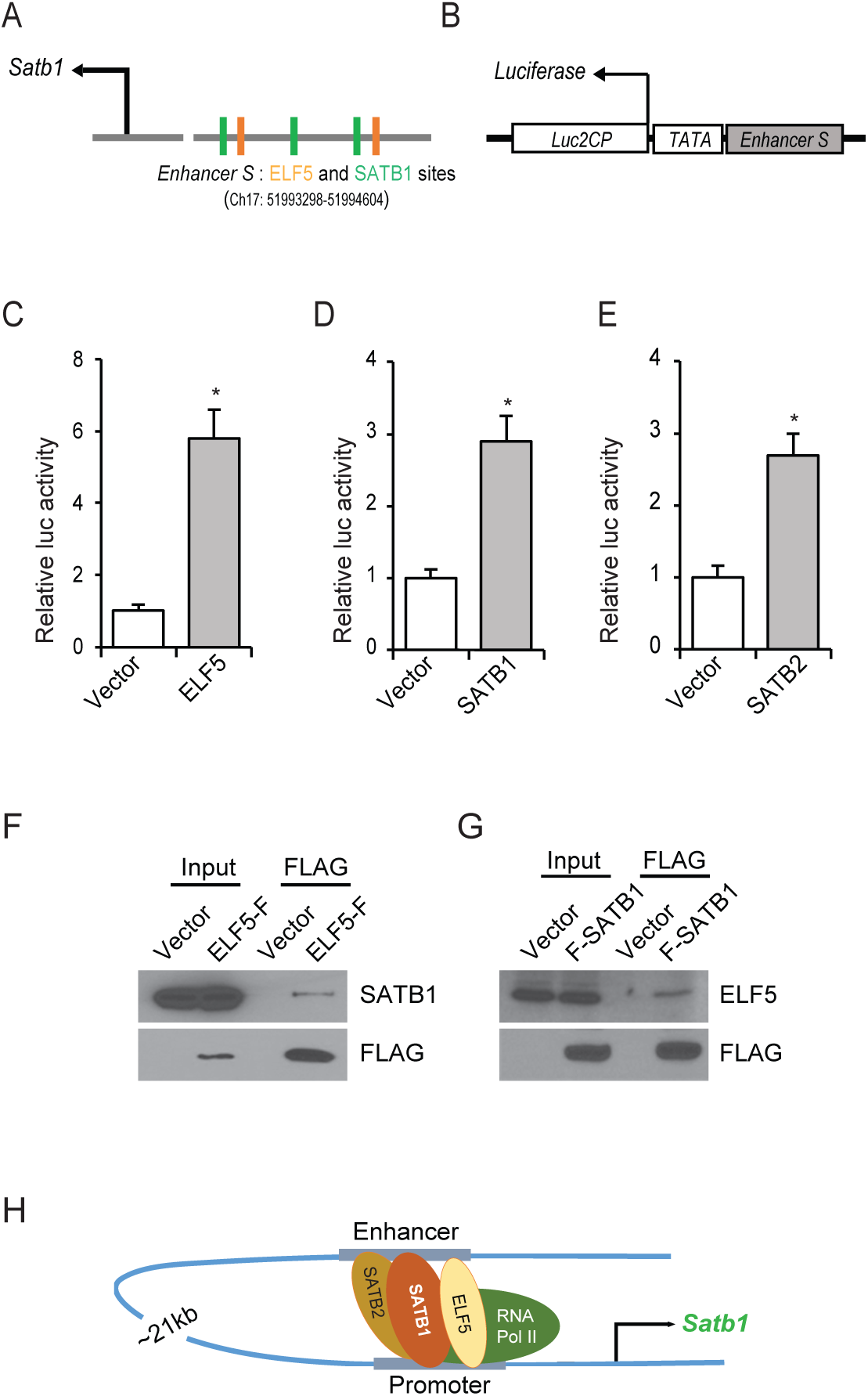
ELF5-SATB1 interaction within the enhancer S. A) Schematic diagram showing the TFBSTools-detected two ELF5 binding sites near SATB1 motifs in mouse *Satb1* enhancer sequence. B) An enhancer-reporter construct was prepared by cloning 1.5 Kb of *enhancer S* upstream of a minimal TATA promoter within the *Luc2CP* firefly luciferase vector. C-E), Ectopic expression of ELF5, SATB1 or SATB2 in Rcho1 rat TS cells significantly upregulated the promoter-reporter activity. Furthermore, co-immunoprecipitation of either ELF5 or SATB1 with Rcho1 nuclear proteins exhibited that SATB1 interacts with ELF5 in trophoblast cells (F, G). Taken together, we propose a model of ELF5-SATB interaction that regulates *Satb1* expression in the trophoblast stem-state (H). Luciferase assay data are expressed as the means ± S.D. *, p< 0.05 (n=3). ELF5-F, ELF5 with C-terminal FLAG tag; F-SATB1, SATB1 with N-terminal FLAG Tag.

## DISCUSSION

SATB proteins play essential regulatory roles in a range of stem cells [15-17, 19]. During early embryonic development, ES, TS, and XEN cells are the three stem cell lineages that give rise to the embryo proper, placenta, and yolk sac, respectively. Among these three stem cell lineages, only TS cells exhibit robust expression of SATB1 (Fig. 1 and S1). However, *Satb1* was induced during reprogramming of mouse ES cells to TS cells, which was also reported in a previous study [26]. Such induction of *Satb1* expression during reprogramming of ES cells to trophoblast fate indicates that trophoblast-specific cell signaling facilitates the expression. It has recently been shown that FGF4 signaling, which is essential for TS cell maintenance, may impact *Satb1* expression in mouse preimplantation embryos [17].

Expression of *Satb1* in trophoblast cells has been reported to be stem-state-specific both *in vivo* and *in vitro* [18, 19]. Differential expression of *Satb*1 in the trophoblast stem-state suggests an important role for stem-specific transcriptional regulators controlling its expression. However, the upstream transcription factors that regulate stem-state specific expression of *Satb1* in trophoblast cells are still unknown.

*Satb1* is an essential regulator of T cell differentiation and FoxP3 plays an important role in transcriptional repression of *Satb1* in regulatory T cells [37]. *Satb1* is also an important chromatin regulator in epidermis, where p63 is essential for maintaining *Satb1* gene expression [38]. However, based on available GEO data (GSE12999 and GSE21938) expression of both FoxP3 and p63 is very low in TS cells, and they do not show any change in expression with induction of differentiation [18, 26]. These findings suggest that regulation of *Satb1* in trophoblast cells is different from T cells and epidermis. To explore the trophoblast-specific *Satb1* regulation, we identified *Satb1* promoters in TS cells. In contrast to T cells that express all four *Satb1* variants, only variant 1 and 2 transcripts were detected in trophoblast cells, with variant 2 being predominant. These proximal promoters were enriched with H3K27ac and H3K4me3, which are marks of active promoters [39]. Presence of CpG islands within the promoters of *Satb1* suggests its potential role as a master developmental regulator [40-42].

An enhancer region ∼21kbp upstream of the *Satb1* variant 2 promoter was identified based on active histone marks [29]. Changes in H3K27ac activity in this enhancer region (*enhancer S*) correlated with *Satb1* expression levels in trophoblast cells. Requirement of the enhancer for *Satb1* expression was demonstrated by CRISPR/CAS9 mediated targeted deletion of this region. Targeted deletion of *enhancer S* reduced *Satb1* expression, which caused differentiation of Rcho1 TS cells maintained in proliferating media. This observation is in line with our previous report that found induction of TS cell differentiation following *Satb1* knockdown [19]. However, trophoblast differentiation due to other reasons can also lead to inhibition of *Satb1* expression. We utilized a transient induction of CRISPR interference to avoid the effect of cell differentiation. CRISPR interference provided direct evidence for the importance of this enhancer in regulating *Satb1* expression in TS cells.

Long-range chromatin interactions can occur intrachromosomally or interchromosomally [43, 44]. Intrachromosomal interactions have been reported between promoters and enhancers located far away from each other [43, 44]. In this study, we detected a chromatin looping of the cis-acting enhancer to the *Satb1* variant 2 promoter across a 21kbp distance. Bioinformatic analyses indicated potential ELF5 binding sites near SATB1 binding sites within *enhancer S* region. ChIP and reporter assays demonstrated that ELF5 and SATB homeobox proteins bind to *enhancer S* and had a stimulatory effect on the enhancer-activity. Binding of ELF5, SATB1 and SATB2 was also detected within the proximal promoter (Fig. S3 E-G). These findings suggest that the looping interaction between the enhancer and the proximal promoter in mouse TS cells was mediated by SATB proteins in association with ELF5. In TS cells, ELF5 can interact with other transcription factors and act as a molecular switch regulating cell differentiation [45]. SATB1 and SATB2 can also form heterodimers to regulate gene expression [19, 46]. It is also well-known that SATB1 can mediate long-range chromatin interactions for gene regulation [4, 47, 48]. Thus, ELF5 interaction with SATB1 to regulate gene expression over a long distance is a plausible mechanism of the transcriptional regulation of *Satb1*.

Trophoblast stem-specific *Satb1* expression suggests that differentially expressed stem-factors may play a crucial role in regulation of *Satb1*. Indeed, SATB proteins as well as ELF5 exhibited trophoblast stem-specific differential expression both in vivo and in vitro (Fig. S4 A-F). We identified that ELF5 plays an important role in regulating *Satb1* expression (Fig. S4 G, H). Developmentally, expression of ELF5 is restricted to the trophoblast lineage and creates a positive feedback loop with other TS cell determinants [49]. We previously demonstrated that SATB proteins contribute to the TS cell stem-state by sustaining the expression of TS factors [19]. Therefore, it is likely that SATB proteins interact with ELF5 in TS cells to augment a positive feedback loop to maintain the trophoblast stem-state.

## ACKNOWLEDGEMENTS

This research was supported by National Institute of Health Grants HD079363 (M.A.R., W.Y., and V.P.C) and HD079545 (G.T.).

## AUTHOR CONTRIBUTIONS

M.A.R. conceived and coordinated the study and prepared the manuscript. W.Y., S.B., V.P.C, A.R., R.R.S. and K.D. performed the experiments and analyzed that data. S.B. prepared the figures and edited the manuscript. M.W.W. and G.T. contributed to designing and editing the manuscript. All the authors approved final version of the manuscript.

---

**Fig. S1. *Satb1* transcript variants and control experiments for ChIP assays.** A) Schematic diagram showing the reference 5’ sequences of four different transcript variants of mouse *Satb1*. B) The accession numbers, noncoding variant specific first exons, common second exons, coding sequences (CDS), and the transcription start sites on mouse chromosome 17 are presented in a tabulated form. Mouse positive and negative control primer sets were used for validating the ChIP assays (Fig. S1). ChIP-qPCR data are expressed as the means ± S.D. *, p< 0.05 (n=3). NC, Negative Control Primer Set; PC, Positive Control Primer Set.

## REFERENCES

1. Dickinson, L.A., et al., A tissue-specific MAR/SAR DNA-binding protein with unusual binding site recognition. Cell, 1992. 70(4): p. 631–45.

2. Dickinson, L.A., C.D. Dickinson, and T. Kohwi-Shigematsu, An atypical homeodomain in SATB1 promotes specific recognition of the key structural element in a matrix attachment region. J Biol Chem, 1997. 272(17): p. 11463–70.

3. Dobreva, G., J. Dambacher, and R. Grosschedl, SUMO modification of a novel MAR-binding protein, SATB2, modulates immunoglobulin mu gene expression. Genes Dev, 2003. 17(24): p. 3048–61.

4. Yasui, D., et al., SATB1 targets chromatin remodelling to regulate genes over long distances. Nature, 2002. 419(6907): p. 641–5.

5. Cai, S., H.J. Han, and T. Kohwi-Shigematsu, Tissue-specific nuclear architecture and gene expression regulated by SATB1. Nat Genet, 2003. 34(1): p. 42–51.

6. Cai, S., C.C. Lee, and T. Kohwi-Shigematsu, SATB1 packages densely looped, transcriptionally active chromatin for coordinated expression of cytokine genes. Nat Genet, 2006. 38(11): p. 1278–88.

7. Alvarez, J.D., et al., The MAR-binding protein SATB1 orchestrates temporal and spatial expression of multiple genes during T-cell development. Genes Dev, 2000. 14(5): p. 521–35.

8. Notani, D., et al., Global regulator SATB1 recruits beta-catenin and regulates T(H)2 differentiation in Wnt-dependent manner. PLoS Biol, 2010. 8(1): p. e1000296.

9. Nakayama, Y., et al., A nuclear targeting determinant for SATB1, a genome organizer in the T cell lineage. Cell Cycle, 2005. 4(8): p. 1099–106.

10. Wen, J., et al., SATB1 family protein expressed during early erythroid differentiation modifies globin gene expression. Blood, 2005. 105(8): p. 3330–9.

11. Dobreva, G., et al., SATB2 is a multifunctional determinant of craniofacial patterning and osteoblast differentiation. Cell, 2006. 125(5): p. 971–86.

12. Alcamo, E.A., et al., Satb2 regulates callosal projection neuron identity in the developing cerebral cortex. Neuron, 2008. 57(3): p. 364–77.

13. Britanova, O., et al., Satb2 is a postmitotic determinant for upper-layer neuron specification in the neocortex. Neuron, 2008. 57(3): p. 378–92.

14. Gyorgy, A.B., et al., SATB2 interacts with chromatin-remodeling molecules in differentiating cortical neurons. European Journal of Neuroscience, 2008. 27(4): p. 865–873.

15. Will, B., et al., Satb1 regulates the self-renewal of hematopoietic stem cells by promoting quiescence and repressing differentiation commitment. Nat Immunol, 2013. 14(5): p. 437–45.

16. Savarese, F., et al., Satb1 and Satb2 regulate embryonic stem cell differentiation and Nanog expression. Genes Dev, 2009. 23(22): p. 2625–38.

17. Goolam, M. and M. Zernicka-Goetz, The chromatin modifier Satb1 regulates cell fate through Fgf signalling in the early mouse embryo. Development, 2017. 144(8): p. 1450–1461.

18. Kent, L.N., T. Konno, and M.J. Soares, Phosphatidylinositol 3 kinase modulation of trophoblast cell differentiation. BMC Dev Biol, 2010. 10: p. 97.

19. Asanoma, K., et al., SATB homeobox proteins regulate trophoblast stem cell renewal and differentiation. J Biol Chem, 2012. 287(3): p. 2257–68.

20. Cockburn, K. and J. Rossant, Making the blastocyst: lessons from the mouse. J Clin Invest, 2010. 120(4): p. 995–1003.

21. Roberts, R.M. and S.J. Fisher, Trophoblast stem cells. Biol Reprod, 2011. 84(3): p. 412–21.

22. Pfeffer, P.L. and D.J. Pearton, Trophoblast development. Reproduction, 2012. 143(3): p. 231–46.

23. Tanaka, S., et al., Promotion of trophoblast stem cell proliferation by FGF4. Science, 1998. 282(5396): p. 2072–5.

24. Kunath, T., et al., Imprinted X-inactivation in extra-embryonic endoderm cell lines from mouse blastocysts. Development, 2005. 132(7): p. 1649–61.

25. Sahgal, N., et al., Rcho-1 trophoblast stem cells: a model system for studying trophoblast cell differentiation. Methods Mol Med, 2006. 121: p. 159–78.

26. Ralston, A., et al., Gata3 regulates trophoblast development downstream of Tead4 and in parallel to Cdx2. Development, 2010. 137(3): p. 395–403.

27. Amita, M., et al., Complete and unidirectional conversion of human embryonic stem cells to trophoblast by BMP4. Proc Natl Acad Sci U S A, 2013. 110(13): p. E1212–21.

28. Untergasser, A., et al., Primer3Plus, an enhanced web interface to Primer3. Nucleic Acids Res, 2007. 35(Web Server issue): p. W71–4.

29. Tuteja, G., T. Chung, and G. Bejerano, Changes in the enhancer landscape during early placental development uncover a trophoblast invasion gene-enhancer network. Placenta, 2016. 37: p. 45–55.

30. Dale, R.K., B.S. Pedersen, and A.R. Quinlan, Pybedtools: a flexible Python library for manipulating genomic datasets and annotations. Bioinformatics, 2011. 27(24): p. 3423–4.

31. Chuong, E.B., et al., Endogenous retroviruses function as species-specific enhancer elements in the placenta. Nat Genet, 2013. 45(3): p. 325–9.

32. Kabadi, A.M. and C.A. Gersbach, Engineering synthetic TALE and CRISPR/Cas9 transcription factors for regulating gene expression. Methods, 2014. 69(2): p. 188–97.

33. Dekker, J., et al., Capturing chromosome conformation. Science, 2002. 295(5558): p. 1306–11.

34. Tan, G. and B. Lenhard, TFBSTools: an R/bioconductor package for transcription factor binding site analysis. Bioinformatics, 2016. 32(10): p. 1555–6.

35. Wenger, A.M., et al., PRISM offers a comprehensive genomic approach to transcription factor function prediction. Genome Res, 2013. 23(5): p. 889–904.

36. Lee, D.S., et al., In vivo genetic manipulation of the rat trophoblast cell lineage using lentiviral vector delivery. Genesis, 2009. 47(7): p. 433–9.

37. Beyer, M., et al., Repression of the genome organizer SATB1 in regulatory T cells is required for suppressive function and inhibition of effector differentiation. Nat Immunol, 2011. 12(9): p. 898–907.

38. Fessing, M.Y., et al., p63 regulates Satb1 to control tissue-specific chromatin remodeling during development of the epidermis. J Cell Biol, 2011. 194(6): p. 825–39.

39. Consortium, E.P., An integrated encyclopedia of DNA elements in the human genome. Nature, 2012. 489(7414): p. 57–74.

40. Ponger, L., L. Duret, and D. Mouchiroud, Determinants of CpG islands: expression in early embryo and isochore structure. Genome Res, 2001. 11(11): p. 1854–60.

41. Tanay, A., et al., Hyperconserved CpG domains underlie Polycomb-binding sites. Proc Natl Acad Sci U S A, 2007. 104(13): p. 5521–6.

42. Vavouri, T. and B. Lehner, Human genes with CpG island promoters have a distinct transcription-associated chromatin organization. Genome Biol, 2012. 13(11): p. R110.

43. Deng, W. and G.A. Blobel, Do chromatin loops provide epigenetic gene expression states? Curr Opin Genet Dev, 2010. 20(5): p. 548–54.

44. Dean, A., In the loop: long range chromatin interactions and gene regulation. Brief Funct Genomics, 2011. 10(1): p. 3–10.

45. Latos, P.A., et al., Elf5-centered transcription factor hub controls trophoblast stem cell self-renewal and differentiation through stoichiometry-sensitive shifts in target gene networks. Genes Dev, 2015. 29(23): p. 2435–48.

46. Zhou, L.Q., et al., The AT-rich DNA-binding protein SATB2 promotes expression and physical association of human (G)gamma- and (A)gamma-globin genes. J Biol Chem, 2012. 287(36): p. 30641–52.

47. Gong, F., et al., The BCL2 gene is regulated by a special AT-rich sequence binding protein 1-mediated long range chromosomal interaction between the promoter and the distal element located within the 3’-UTR. Nucleic Acids Res, 2011. 39(11): p. 4640–52.

48. Yang, Y., et al., SATB1 Mediates Long-Range Chromatin Interactions: A Dual Regulator of Anti-Apoptotic BCL2 and Pro-Apoptotic NOXA Genes. PLoS One, 2015. 10(9): p. e0139170.

49. Ng, R.K., et al., Epigenetic restriction of embryonic cell lineage fate by methylation of Elf5. Nat Cell Biol, 2008. 10(11): p. 1280–90.

