## Supplemental Table S1-S8 for "A long-range chromatin interaction regulates SATB homeobox 1 gene expression in trophoblast stem cells"

**SUPPLEMENTAL TABLES**

**Table S1. Primers used for conventional RT-PCR analyses**

| Target | Accession no. | Forward primer | Reverse primer | Amplicon (bp) |
|---|---|---|---|---|
| *Mouse Satb1* | NM_009122.2 | CTTTGGAGCAGCAAGTTTCC | AGGTTCTCCCACAGGGTCCT | 568 |
| *Mouse Satb2* | NM_139146.2 | TACTCCAATCCGAAACCAAG | GTTGTCGGTGTCGAGGTTTT | 573 |
| *Mouse Elf5* | NM_001145813.1 | GCTTGAAAACAAGTGGCATC | GTCCGGTGTCCATCAGAGTT | 353 |
| *Mouse Gapdh* | NM_008084.3 | ACCACAGTCCATGCCATCAC | TCCACCACCCTGTTGCTGTA | 452 |

**Table S2. Primers used for conventional RT-PCR and RT-qPCR analyses of mouse *Satb1* transcript variants**

| Target | Accession no. | Forward primer | Reverse primer | Amplicon (bp) |
|---|---|---|---|---|
| *Mouse Var1* | NM_001163630.1 | CCTTCAGGTCTGCTGCTTTT | CACTCCCTGCATCTTTCCAC | 351 |
| *Mouse Var2* | NM_009122.2 | AAGTCAAGTTGCCATTAACTTCC | CACTCCCTGCATCTTTCCAC | 366 |
| *Mouse Var3* | NM_001163631.1 | GCTAACCTGCCAGAGAGAACTT | CACTCCCTGCATCTTTCCAC | 292 |
| *Mouse Var4* | NM_001163632.1 | CCCAGGCAAACAACGACAA | CACTCCCTGCATCTTTCCAC | 262 |
| *Mouse Rn18s* | NR_003278.3 | GCAATTATTCCCCATGAACG | GGCCTCACTAAACCATCCAA | 123 |

**Table S3. Primers used for qRT-PCR analyses of gene expression**

| Symbol | Accession no. | Forward primer | Reverse primer | Amplicon (bp) |
|---|---|---|---|---|
| *Human SATB1* | NM_001131010.3 | GCAGAATTTGTGCTGGTGAG | CCAACCTGGATTAGCCCTTT | 119 |
| *Mouse Satb1* | NM_009122.2 | GTGGCAGACATGCTTCAAGA | TACTGTGGTGTGCGACCATT | 111 |
| *Rat Satb1* | NM_001012129 | TGAGAGGGAAAGGAGCTTGA | TGTTCTCTGGCTTCCCATTC | 131 |
| *Rat Satb2* | NM_001109306 | CTTCCTCAACCTGCCTGAAG | GTTGTCGGTGTCGAGGTTTT | 150 |
| *Rat Cdx2* | NM_023963.1 | CAGGAGGAAAGCTGAGTTGG | TGCTGCTGTTGCAACTTCTT | 120 |
| *Rat Eomes* | XM_017596193.1 | CAATGTGTTCGTGGAAGTGG | GTTGGGAGATTCTGGGTGAA | 133 |
| *Rat Prl3d1* | NM_017363.3 | TCTTCCGGGAGCTTCTGTTA | GACCAGGCAGGGTAGTCAAA | 150 |
| *I8SrRNA/ Rn18s (Human, Mouse and Rat)* | NR_145819.1<br>NR_003278.3<br>NR_046237.1 | GCAATTATTCCCCATGAACG | GGCCTCACTAAACCATCCAA | 123 |

**Table S4. Primers used for ChIP qPCR analyses**

| Target | Forward primer | Reverse primer | Amplicon (bp) |
|---|---|---|---|
| *Mouse Enhancer S* | AGCAGGTGTGAGCAGCTGAG | GACTGTCCTTCAAGTCTTTCTCA | 137 |
| *Rat Enhancer S* | AGCAGGTGTGAGCAGCTGAG | CCTGTCCTTCAAGTCTTTCTCA | 135 |
| *Mouse Var 2 prom* | GGCCACTGAGAAGTTTGGAT | TGAGTGAGTCCCGCTTCTTT | 113 |
| *Rat Var 2 prom* | GGCTCCCGGGATAGAGAAGTT | GGGTGGAGTTGGAAAGGTC | 161 |

Enhancer S, *Satb1* enhancer; Var 2 prom, *Satb1* variant 2 promoter.

**Table S5. gRNAs used for CRISPR- interference and deletion of *enhancer S* and the *Satb1* variant 2 promoter**

| Target | Sites | Forward oligos | Reverse oligos |
|---|---|---|---|
| *Rat Enhancer S* | *1* | rSatb1*Enhancer S* gRNA-1F<br>CACC GAGAGGCACATCCGGTAAGT | rSatb1 *Enhancer S* gRNA-1R<br>AAAC ACTTACCGGATGTGCCTCTC |
| *Rat Enhancer S* | *2* | rSatb1 *Enhancer S* gRNA-2F<br>CACC GCCCTGCAGAACGCACTCTA | rSatb1 *Enhancer S* gRNA-2R<br>AAAC TAGAGTGCGTTCTGCAGGGC |
| *Rat Enhancer S* | *3* | rSatb1 *Enhancer S* gRNA-3F<br>CACC GTGGCGAACAGGTGTCTACT | rSatb1 *Enhancer S* gRNA-3R<br>AAAC AGTAGACACCTGTTCGCCAC |
| *Rat Var 2 prom* | *1* | rSatb1 Var2 pro gRNA-1F<br>CACC CATCCGAAGTGGGCGTTTAA | rSatb1 Var2 pro gRNA-1R<br>AAAC TTAAACGCCCACTTCGGATG |
| *Rat Var 2 prom* | *2* | rSatb1 Var2 pro gRNA-2F<br>CACC GAGCGCGGCGAGCAGCGAGC | rSatb1 Var2 pro gRNA-2R<br>AAAC GCTCGCTGCTCGCCGCGCTC |
| *Rat Var 2 prom* | *3* | rSatb1 Var2 pro gRNA-3F<br>CACC GTAGCGCGCGGCCGAGGGGA | rSatb1 Var2 pro gRNA-3R<br>AAAC TCCCCTCGGCCGCGCGCTAC |

**Color code:** Blue- cloning sites; Black: target sequence; rSatb1, rat *Satb1*; Var 2 prom, Variant 2 promoter

**Table S6. Primers used for conventional PCR to detect CRISPR/Cas9 mediated enhancer deletion**

| Target | Forward primer | Reverse primer | Amplicon (bp) |
|---|---|---|---|
| *Rat Enhancer S* | ACTTACCGGATGTGCCTCTC | TAGAGTGCGTTCTGCAGGGC | 506 |
| *Control site (Var 2 prom)* | GGCTCCCGGGATAGAGAAGTT | GGGTGGAGTTGGAAAGGTC | 161 |

**Table S7. PCR Primers used for Chromatin Conformation Capture (3C) analyses**

| **Primer** | **Target** | **Location** | **Target sequence** | **Amplicon (bp)** |
|---|---|---|---|---|
| -20298Rv | *Mouse Enhancer S* | Chr17: 51992270<br>506 bp from *Bgl* II site | GAGCTGAATGAGCCTGAAAGA | 647 |
| +486Rv | Mouse var 2 promoter | Chr17: 51971486<br>141 bp from *Bgl* II site | TGGCCCGCTTTAGAGGACGAAG | |
| -20020Rv | *Enhancer S* sequencing | Chr17: 51991992 | GTTCAATTGGCCAGGGTTATTT | |
| Control-F | Enhancer | | AGCAGGTGTGAGCAGCTGAG | 137 |
| Control-R | Enhancer | | GACTGTCCTTCAAGTCTTTCTCA | |

**Table S8. shRNA target site and sequences**

| **shRNA** | **Target** | **Target sequence** | **shRNA sequence** (cloned into *Age*I and *Ecor*I sites of pLKO.1) |
|---|---|---|---|
| *shElf5.2* | Rat *Elf5* | ATCAGATCAAACTAGACATTT | accggtATCAGATCAAACTAGACATTTctcgagAAATGTCTAGTTTGATCTGATTTTTTTGgaattc |
| *shElf5.4* | Rat *Elf5* | GCCCTGAGATACTACTATAAA | accggtGCCCTGAGATACTACTATAAActcgagTTTATAGTAGTATCTCAGGGCTTTTTTgaattc |
| *shSCR* | None | CCTAAGGTTAAGTCGCCCTC | accggtCCTAAGGTTAAGTCGCCCTCGctcgagCGAGGGCGACTTAACCTTAGGTTTTTTgaattc |

**Color code:** Blue- cloning sites; Black: sense target; Orange: shRNA loop; Green: antisense target; Red: stop signal for RNA polymerase III
