## Supplemental Figure 1 for "A long-range chromatin interaction regulates SATB homeobox 1 gene expression in trophoblast stem cells"

Figure S1

A

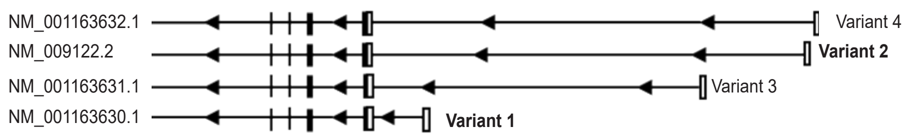


B

| Variant | Reference mRNA | Exon 1 | Exon 2 | CDS | Transcription start site |
|---|---|---|---|---|---|
| 1 | NM_001163630.1 | 1-287 | 288-532 | 322-2616 | Chr17: 51951379 |
| 2 | NM_009122.2 | 1-178 | 179-423 | 213-2507 | Chr17: 51971972 |
| 3 | NM_001163631.1 | 1-155 | 156-400 | 190-2484 | Chr17: 51965913 |
| 4 | NM_001163632.1 | 1-128 | 129-373 | 163-2457 | Chr17: 51972615 |

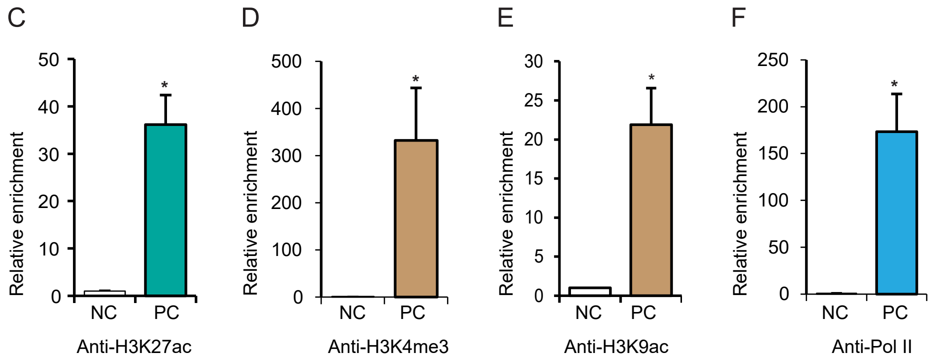
